# Incorporating motor preparation time transforms micro-offline gains into micro-offline losses

**DOI:** 10.64898/2026.04.25.720821

**Authors:** NA Ahmed, T. Suresh, SJ Hussain, M. Freedberg

**Affiliations:** The University of Texas at Austin, Department of Kinesiology and Health Education; The University of Iowa, Department of Health, Sport, and Human Physiology, Department of Neurology; The University of Iowa, Department of Psychiatry

**Keywords:** motor learning, micro-offline gains, micro-consolidation, offline learning

## Abstract

During explicit sequence learning (ESL), micro-offline gains (MOGS) occur during brief rest periods. MOGS are calculated as the difference in keypresses-per-second (KPS) between the first sequence of one trial and the last sequence of the preceding trial. To date, all studies evaluating MOGS have calculated KPS from the motor execution time (MET) that occurs between keypresses, but this approach ignores potential contributions from motor preparation which occur prior to the first keypress. Given that ESL relies on both pre-movement motor planning and subsequent motor execution, we hypothesized that ignoring motor preparation time (MPT) neglects a critical component of skill acquisition, potentially misrepresenting the true magnitude of MOGS. To test this, we calculated MOGS with and without MPT in thirty adults who performed an ESL task. Our results show that including MPT flipped MOGS from positive to negative and significantly increased the positive correlation between early learning and a gold-standard ESL metric: the number of correct sequences performed. Our results suggest that MPT should be incorporated into MOGS calculations and that excluding it overestimates micro-offline learning.

## Introduction

Motor skill learning involves acquiring and refining movements through practice and is a critical component of daily life. Motor skill acquisition refers to the initial encoding of novel movement patterns through practice, resulting in improvements in effector coordination and performance. Following skill acquisition, motor skill memories undergo consolidation, which is a time-dependent offline process that stabilizes these newly formed fragile memory traces into robust long-term memories (Walker et al., 2002; Yang et al., 2025). Consolidation supports long-term skill retention and can be measured as resistance to interference or performance gains that occur over either wakeful rest or sleep (Albouy et al., 2015; Krakauer & Shadmehr, 2006; Robertson, 2012).

Motor skill learning relies on a network of sensorimotor regions. Individuals with damage to this network, including primary motor cortex (M1), often need to relearn everyday motor skills (Kogan et al., 2023; Krakauer & Mazzoni, 2011; Nguyen et al., 2025; Nudo, 2013; Nudo et al., 1996). To facilitate motor skill relearning, neuromotor rehabilitation specialists rely on results of research using scientifically validated motor learning tasks and metrics. One commonly used experimental paradigm for measuring motor skill acquisition and consolidation is the explicit sequence learning (ESL) task (Karni et al., 1998). The ESL task requires learners to perform a sequence of keypresses, similar to playing a melody on a piano, as quickly and accurately as possible. ESL acquisition is typically measured as the increase in the number of correct sequences from the first to last trials of an acquisition period, while consolidation is often measured as the difference in number of correct sequences between the last trials of an acquisition period and all trials of a subsequent retention period (Robertson et al., 2004; Walker et al., 2002). The number of correct sequences is a validated metric of motor learning (Fischer et al., 2002; Herszage et al., 2021; Hsu et al., 2023; Karni et al., 1998; Walker et al., 2002), which accurately represents ESL and is often used to quantify both online ESL gains that accumulate during the acquisition phase and offline ESL gains that occur during consolidation (Fischer et al., 2002; Herszage et al., 2021; Karni et al., 1998; Walker et al., 2002). Because the ESL task involves conscious sequence awareness, rapid performance improvements early during acquisition are primarily driven by improvements in cognitive control over sequence execution (Robertson, 2007).

Prior work has repeatedly shown that ESL consolidation occurs over both wakeful rest lasting from minutes to hours (Wang et al., 2024) and extended periods of sleep (Albouy et al., 2008). However, recent studies suggest that ESL skill consolidation can also occur during short inter-trial rest periods lasting only 10 seconds (Bönstrup et al., 2019; Jacobacci et al., 2020). These improvements are referred to as “micro-offline gains” (MOGS) and are calculated by measuring changes in key tapping speed between the last correct sequence of one trial and the first correct sequence of the next trial. Using this approach, positive values indicate MOGS. Furthermore, online ESL gains, referred to as “micro-online gains” (MOnGS), are calculated as the difference in tapping speed between the first and last correct sequences within a trial. Learning is thus quantified as the sum of MOGS and MOnGS during early learning, with early learning defined as the period during which 95% of performance improvements occur. The original MOGS report (Bönstrup et al., 2019) and several others since (Buch et al., 2021; Jacobacci et al., 2020; Mylonas et al., 2024) have defined key tapping speed as keypresses per second (KPS), which is obtained by dividing the number of keypress transitions by the total elapsed time between the first and last keypresses. However, to our knowledge, neither the original report or any studies since have systematically validated this metric.

A comprehensive ESL metric should accurately characterize all or most learning-related improvements in performance. Each ESL trial includes two phases. First, motor preparation time (MPT) represents the time from the ‘go’ cue to the first keypress. Second, movement execution time (MET) is the time between subsequent keypresses (Abrahamse et al., 2013; Brown et al., 2022; Diedrichsen & Kornysheva, 2015; Haith et al., 2015). Critically, successful ESL performance requires both motor preparation and motor execution (Ariani et al., 2020), and because MPT decreases as motor learning occurs, learning involves both MPT and MET. Yet, most studies calculate MOGS using only MET (Bönstrup et al., 2019, 2020; Gann et al., 2023; Jacobacci et al., 2020; Mylonas et al., 2024), ignoring potential contributions from MPT. This omission systematically overestimates the speed (KPS) of the first post-rest sequence by discounting the movement planning time prior to the first keypress (Ariani & Diedrichsen, 2019; Bashford et al., 2022; Brown et al., 2022; Diedrichsen & Kornysheva, 2015). Because movement planning time is longest during early learning (Ariani et al., 2020; Ariani & Diedrichsen, 2019; Bashford et al., 2022), excluding MPT likely inflates MOGS. Here, we propose that MOGS measurements that capture the *total time* required to both plan and execute a sequence better captures ESL.

In this study, we systematically compared MOGS measurements obtained during ESL using two distinct methods of quantifying keypresses-per-second: the commonly used method that only captures motor execution gains (MOGS-MET) and a novel metric that incorporates both MPT and MET (MOGS-TOTAL). Here, we tested the hypothesis that including MPT in MOGS would significantly change its value while also strengthening the relationship between early learning (MOGS+MOnGS) and and the gold-standard of ESL: the number of correct sequences completed.

## Methods

### Experimental design and procedure

We recruited thirty right-handed adults (13 male, 17 female) with a mean age of 18.9 years (SD = 1.09). During ESL trials, the sequence presented on the computer screen was 4-1-3-2-4, with the numbers 1-4 corresponding to the index, middle, ring, and pinky fingers, respectively (**Fig. 1A**). Participants were instructed to accurately complete as many sequences as possible in the 10-second trial periods using their non-dominant left hand. Each keypress response added a yellow asterisk to the screen just below the presented sequence. This response feedback was identical to that used in the original report of MOGS by Bönstrup et al. (2019) and was included to confirm that each keypress was registered by the E-prime software.

**Figure 1.**
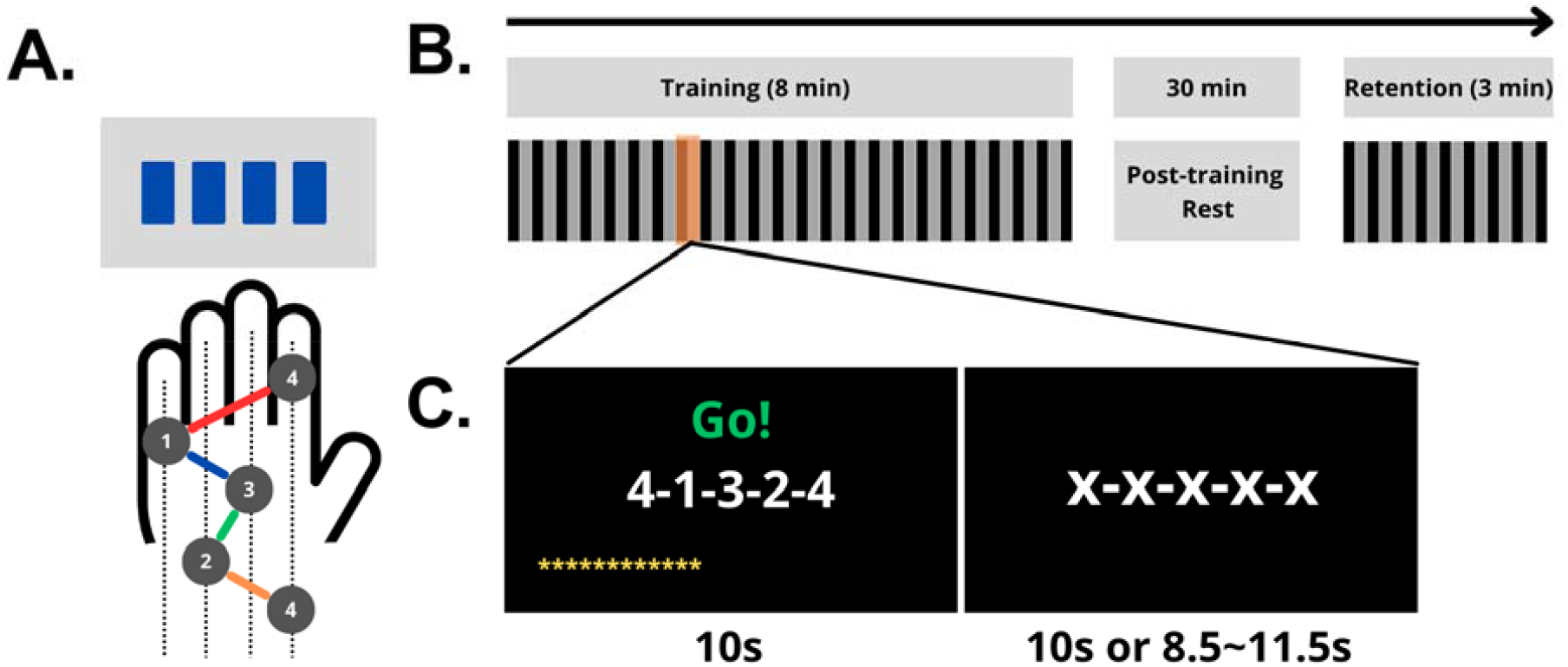
Experimental procedures and tasks. (A) During each ESL trial, participants repeatedly typed the 4-1-3-2-4 sequence on a response pad using their left index, pinky, middle, ring, and index fingers, respectively. (B) Participants completed 24 trials of the ESL task during the acquisition period and 9 trials during the retention period. Each trial (acquisition, retention) included periods of active typing (black lines) and inter-trial rest (grey lines). (C) Each practice trial lasted 10s, followed by rest intervals that were either fixed at 10s (no-jitter group) or jittered between 8.5–11.5s (jitter group).

The data reported here were obtained as part of a larger study evaluating the effects of pre-trial temporal predictability on MOGS and MOnGS. Participants were thus divided into two groups: a jitter group (N=15, 9 male, 6 female, mean age 18.7 ± 1.1 years) and a no-jitter group (N=15, 4 male, 11 female, mean age = 19 ± 1.1 years). In the jitter group, 15 participants completed the task with jittered inter-trial intervals that ranged from 8.5 to 11.5 seconds (mean = 10 seconds). In the no-jitter group, participants completed the task with fixed 10-second rest intervals. Because we observed no significant difference in either MOGS-MPT or MOGS-TOTAL between groups (see *Results*), data were pooled across groups prior to analysis.

Procedures for the experiment are shown in **Figure 1B**. Participants were instructed that when the task begins, they will see a sequence of numbers on the computer monitor and they should complete as many accurate trials as possible with their non-dominant hand until the 10 second trial ended. The sequence was displayed in the center of a computer monitor, and responses were recorded using a serial response box controlled via E-Prime (Psychology Software Tools, Inc., 2016). Participants were not informed of which group they were assigned to and were instructed to rest quietly during the inter-trial rest periods while maintaining their finger position atop the response box. During inter-trial rest periods, the sequence of numbers shown on the screen was replaced by a series of X’s (**Fig. 1C**) as in Bönstrup et al. (2019). After completing 24 ESL trials during the acquisition period, participants were instructed to rest for 30 minutes. During this 30-minute rest, participants remained seated at the experimental setup and were instructed to stay awake, rest quietly, and refrain from practicing the motor task. Then, all participants completed an additional 9 ESL trials, with all of these trials including a fixed 10-second inter-trial rest period.

### Data analysis

Within each 10-second ESL trial, a single sequence was considered correct when all elements of the 4-1-3-2-4 sequence were performed accurately in succession. If the first sequence of a trial was incorrect, we used the next accurately performed sequence. For the first correct 4-1-3-2-4 sequence of each trial, we extracted 1) MPT, which is the latency from the “go” cue at the start of each trial to the first keypress, and 2) MET, which is the latency from the first keypress to the last keypress in a correct sequence (the time between the first ‘4’ and the final ‘4’ in the 4-1-2-3-4 sequence). For the last sequence in a trial, only MET was used to calculate KPS, since there was no MPT involved.

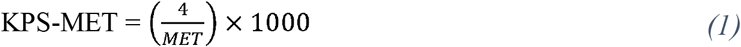

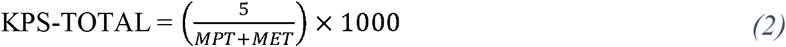

Using MPT and MET, we calculated KPS in two ways: 1) KPS-MET, which only includes MET in KPS (Eq. 1), similar to prior reports, and 2) KPS-TOTAL, which is a novel metric that includes both MPT and MET in KPS (Eq. 2). For both metrics, MOGS was calculated by subtracting the KPS of the last correct sequence of one trial from that of the first correct sequence of the following trial. We performed similar calculations for MOnGS by subtracting the tapping speed of the first correct sequence from the tapping speed of the last correct sequence in the same trial.

Using both methods of calculating MOGS and MOnGS, we calculated early learning as the cumulative sum of all MOGS and MOnGS across the first 11 trials during the acquisition period. As in Bönstrup et al. (2019), we defined the early learning phase as trials 1-11. By trial 11, participants performance reached 90% of their maximum acquisition phase performance (**Fig. 2A**). This approach ensures that our analysis remained entirely within the early learning phase and is consistent with prior reports (Bönstrup et al., 2019).

**Figure 2.**
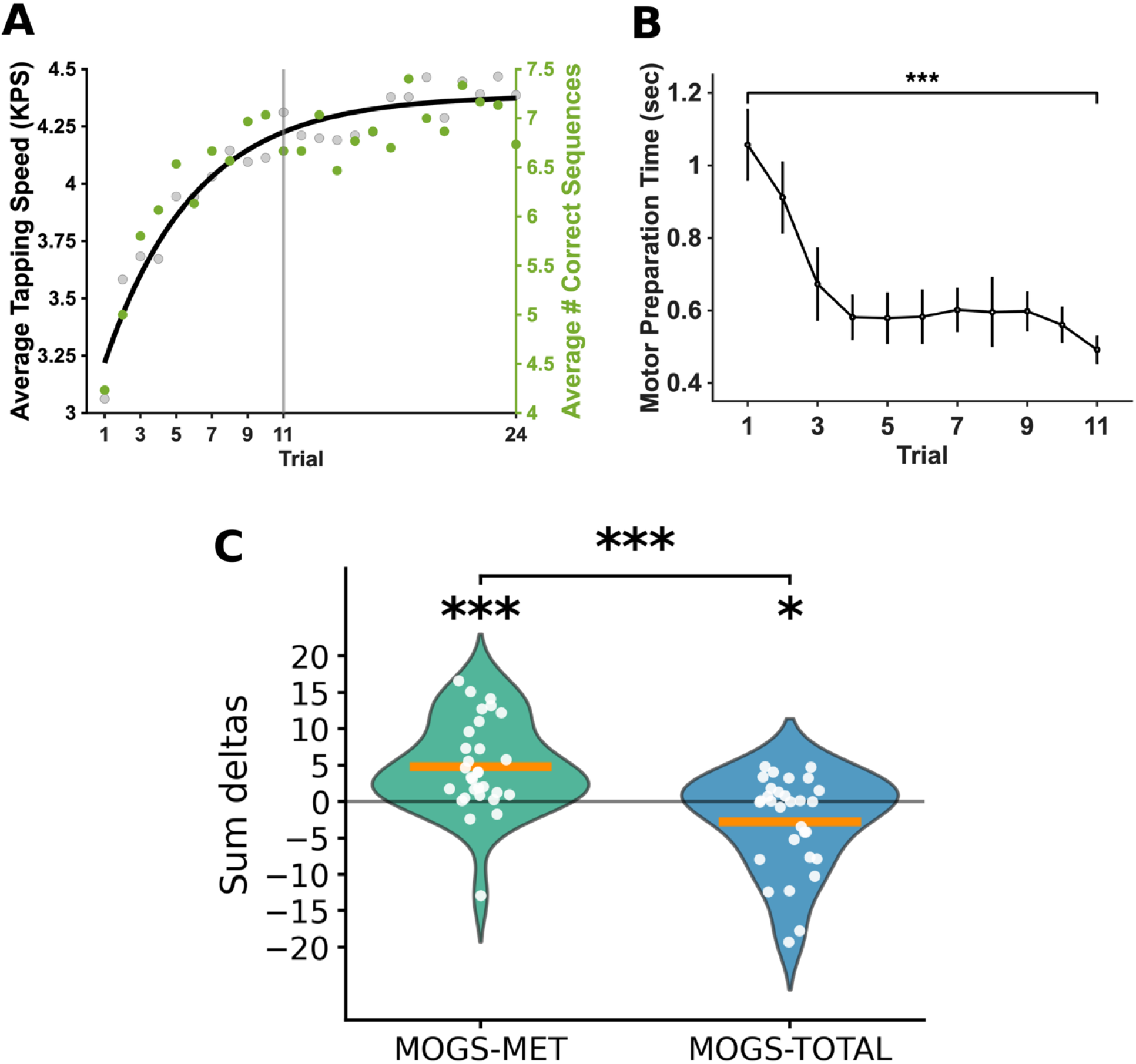
Learning curve, motor preparation time and micro-offline gains. (A) Learning curve. Gray dots show the average tapping speed (KPS) within correct sequences per trial for all participants, as plotted on the left axis. Green dots show the mean number of correct sequences performed per trial for all participants, as plotted on the right axis. The solid black line represents the fitted learning curve for average tapping speed. The vertical line at trial 11 marks the end of the early learning phase, by which point participants achieved 90% of their maximal performance gains. (B) Motor preparation time decreases over early learning trials. Error bars indicate the standard error of the mean. (C) Including MPT in MOGS flips it negative. White dots represent each participant’s MOGS score. Orange bars represent the mean. ***-p□<□.001, *-p□<□.05.

We next calculated the total number of correct sequences performed by the end of early learning. This metric is a simple count of the total number of correct tapping sequences performed without any errors during the first 11 trials. The number of correct sequences is a well-established metric of ESL that has been used for over 30 years; we therefore consider this to be the gold-standard for quantifying ESL. Since all trials have a fixed duration, an increase in the number of completed sequences mathematically necessitates a reduction in the time spent planning (MPT) and/or executing (MET) the movements. Therefore, the number of correct sequences comprehensively captures the total cumulative index of motor learning, integrating improvements in the speed of movement initiation and inter-key transitions.

Retention scores were measured for each participant by subtracting the mean number of correct sequences performed during the last six acquisition trials from the mean number of correct sequences performed during all nine retention trials. To clarify the distinction between motor execution and motor preparation variables, Table 1 provides a comprehensive overview of all metrics and their corresponding calculations.

**Table 1:** Measures and Calculations of Motor Learning Metrics.

| Metric | What it Measures | How it is Calculated |
| --- | --- | --- |
| *Temporal Variables* |  |  |
| MPT | Motor preparation time | Time from "go" cue to 1st keypress |
| MET | Motor execution time | Time from 1st keypress to last keypress |
| *MET Metrics* |  |  |
| KPS-MET | Sequence execution speed (MET only) | $[4 \div \text{MET}] \times 1000$ |
| MOGS-MET | Rest-based gains (MET only) | KPS-MET (1st seq., Trial N) – KPS-MET (last seq., Trial N-1) |
| MOnGS-MET | Within-trial practice gains (MET only) | KPS-MET (last seq.) – KPS-MET (1st seq.) within the same trial |
| Early Learning-MET | Cumulative early skill acquisition (MET only) | Sum of first 11 MOnGS-MET + first 10 MOGS-MET |
| *Comprehensive Metrics (TOTAL)* |  |  |
| KPS-TOTAL | Sequence execution speed (MPT + MET) | $[5 \div (\text{MPT} + \text{MET})] \times 1000$ |
| MOGS-TOTAL | Rest-based gains (MPT + MET) | KPS-TOTAL (first seq., Trial N) – KPS-TOTAL (last seq., Trial N-1) |
| MOnGS-TOTAL | Within-trial practice gains (MPT + MET) | KPS-TOTAL (last seq.) – KPS-TOTAL (1st seq.) within the same trial |
| Early Learning-TOTAL | Cumulative early skill acquisition (MPT + MET) | Sum of first 11 MOnGS-TOTAL + first 10 MOGS-TOTAL |
| *Gold-standard Metric* |  |  |
| Number of correct sequences | Overall learning | Total count of correct sequences across early learning trials |

### Statistical analysis

We first tested whether MPT decreases across ESL trials during the acquisition period, consistent with prior reports (Ariani et al., 2020) using a repeated measures ANOVA with TRIAL as a factor. Second, we asked whether these MPT decreases were associated with MOGS-MET using a Pearson correlation between each participant’s MPT change from trial 1-11 and their mean MOGS-MET. Third, to establish the presence and direction of MOGS-MET and MOGS-TOTAL, we tested both metrics against zero using one-sample t-tests. Fourth, to determine if including MPT in the calculation of MOGS significantly alters the magnitude of calculated MOGS, we conducted paired two-tailed t-tests contrasting MOGS-MET vs MOGS-TOTAL. Fifth, we tested the relationship between the two metrics for calculating early learning with the number of correct sequences performed using three Pearson’s correlations: 1) Early Learning-MET vs. number of correct sequences, 2) Early Learning-TOTAL vs. number of correct sequences, and 3) motor preparation time (MPT) vs. number of correct sequences. Finally, to test our prediction that Early Learning-TOTAL would more positively correlate with the number of correct sequences than Early Learning-MET, we used Steiger’s Z-test to statistically compare the strength of these two correlations. All data were normally distributed (Shapiro-Wilk test), and all analyses were performed in MATLAB 2025b. Alpha was set to 0.05.

## Results

### Between groups analysis (jitter vs. no-jitter groups)

A one-way ANOVA revealed no significant effect of group on the magnitude of MOGS-MET (no-jitter group: mean = 2.8 ± 0.97; jitter group: mean = 4.2 ± 0.98; *F*(1, 28) = 1.11, *p* = .3) or MOGS-TOTAL (fixed-interval: mean = -1.9 ± 1.13; jittered-interval: mean = -2.01 ± 1.2; *F*(1, 28) = 0.005, *p* = .9). Furthermore, one-sample t-tests against zero confirmed that significant MOGS-MET occurred in both groups (no-jitter: *t*(14) = 2.93, *p* = .011; jitter: *t*(14) = 4.41, *p* < .001). We observed no difference in MPT (*t*(28) = 0.68, *p* = .5) and MET (*t*(28) = 1.03, *p* = .3) between the groups.

### MPT as a component of motor learning and its relationship with MOGS

We observed a significant main effect of trial on MPT (*F*(10, 260) = 12.97, *p* < .001) and on MET (*F*(10, 260) = 15.29, *p* < .001), and a post-hoc analyses showed systematic decreases of MPT (F(1, 26) = 28.88, p < .001) and MET (F(1, 26) = 24.11, p < .001) with trials. As illustrated in **Figure 2B**, MPT systematically decreased with trials, showing that improvements in both MET and MPT make up explicit sequence learning. However, we observed no significant relationship between MPT and MOGS-MET (*r*(28) = 0.1, *p* = .5). Thus, our results are consistent with the notion that MPT and MET measure different underlying mechanisms.

### Including MPT in KPS measurements flipped MOGS from positive to negative

Consistent with prior reports, a one-sample t-test confirmed that MOGS-MET are significant and positive (mean = 3.77 ± 0.68), with the mean being significantly greater than zero (*t*(29) = 5.18, *p* < .001; **Fig. 2C**). MOGS-TOTAL, however (mean = -1.96 ± 0.81), was significantly smaller than zero(*t*(29) = -2.41, *p* = .011; **Fig. 2C**), and significantly smaller than MOGS-MET (*t*(29) = 7.4, *p* < .001).

### Relationship to number of correct sequences

A Pearson correlation between Early Learning-MET and number of correct sequences performed was not significant (*r*(28) = 0.2, *p* = .32; **Fig. 3A**). In contrast, Early Learning-TOTAL was significantly and positively correlated with the number of correct sequences performed (*r*(28) = 0.5, *p* = .013; **Fig. 3B**). To determine whether adding MPT to KPS significantly increased the positive relationship between early learning and number of correct sequences, a Steiger’s Z-test for dependent correlations was performed (**Fig. 3C**). This test confirmed our prediction that Early Learning-TOTAL was significantly more positively associated with number of correct sequences performed than Early Learning-MET: *z* = 5.68, *p* < .001, N = 30. Unsurprisingly, a Pearson correlation between MPT and number of correct sequences showed a significant negative correlation (*r*(28) = -0.6, *p* < .001; **Fig. 3D**).

**Figure 3.**
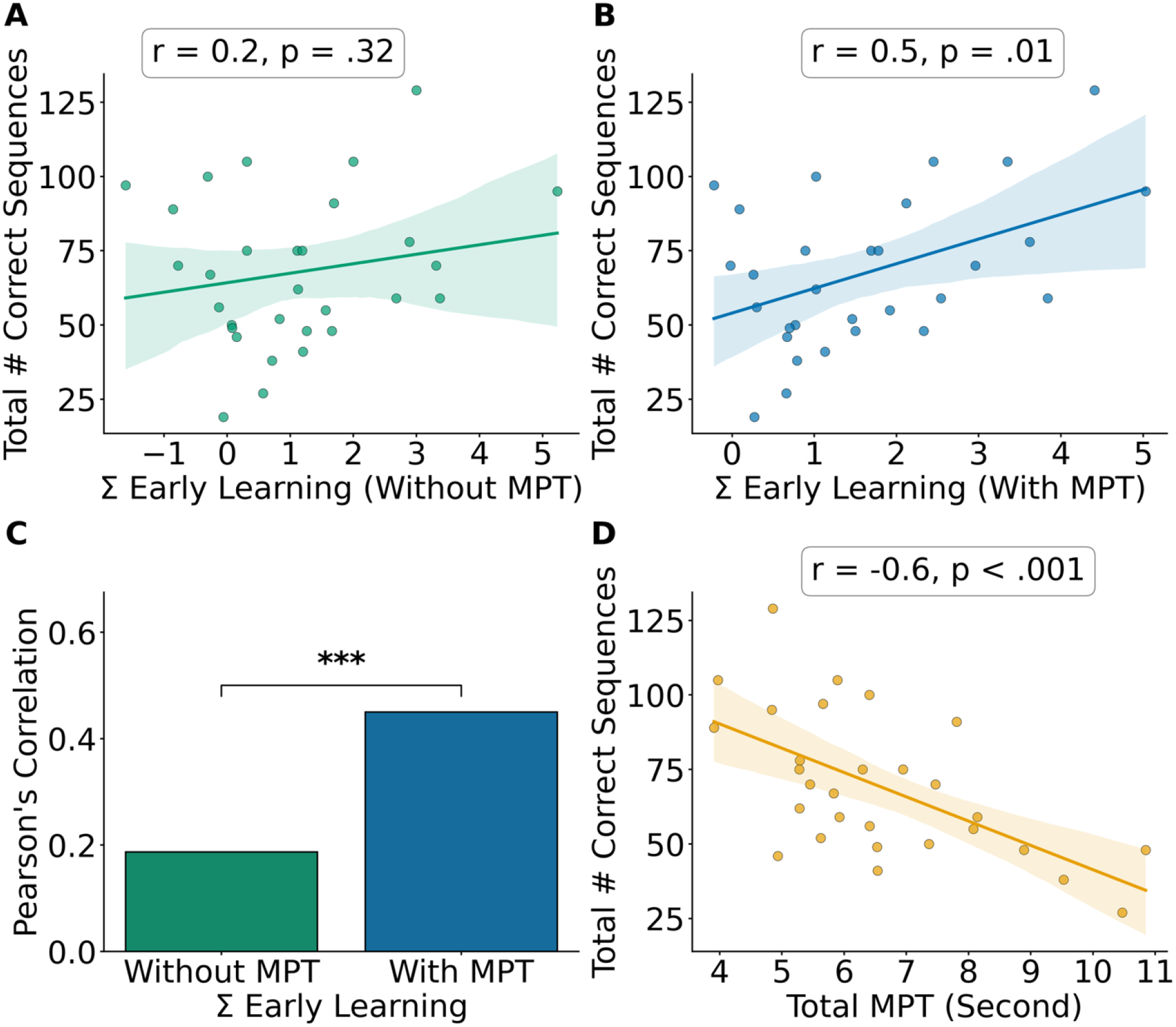
Correlations between Early Learning and number correct sequences. (A) Pearson correlation between early learning calculated without MPT (Early Learning-MET) and number of correct sequences. (B) Pearson correlation between early learning calculated with MPT (Early Learning-TOTAL) and number of correct sequences. (C) Steiger’s Z-test comparing the correlation coefficients from Panels A and B. (D) Pearson correlation between total MPT and number of correct sequences, showing a significant negative relationship where longer planning times predicted lower performance by the end of early learning. In all scatter plots (A, B, D), dots represent individual participant data points, solid lines indicate the linear regression slope, and shaded regions represent the standard error. ***-p□<□.001

### Relationship to retention

Average retention score across all participants was 1.73 ± 0.18 (mean ± SE). Pearson correlations between Early Learning-MET and retention (*r*(28) = -0.3, *p* = .11), and Early Learning-TOTAL and retention (*r*(28) = -0.1, *p* = .61), were not significant.

## Discussion

MOGS represent a new way of measuring motor learning as well as a potential target for interventions aimed at improving function in individuals with sensorimotor network damage. This metric is measured as changes in tapping speed between the last and first sequence performance on consecutive trials. Although this is a new and exciting field of research, the traditional method for calculating MOGS (KPS) has not been rigorously validated and ignores motor preparation time (MPT). We reasoned that, because improvements in MPT are a critical part of motor learning (Ariani et al., 2020; Ariani & Diedrichsen, 2019; Bashford et al., 2022; Brown et al., 2022; Diedrichsen & Kornysheva, 2015), including MPT in MOGS calculations would provide a more comprehensive metric of motor learning than the commonly-used method that focuses solely on MET.

Our study reports several notable findings. First, we confirmed that MPT decreases with practice, highlighting its importance in the motor learning process (Ariani et al., 2020; Ariani & Diedrichsen, 2019; Bashford et al., 2022). These decreases were not associated with MOGS-MET, suggesting that MPT and MET capture independent learning mechanisms. Second, including MPT when calculating MOGS flipped it from positive to negative, showing that not only is MPT important to the calculation of MOGS, but that ignoring it may grossly exaggerate the magnitude of MOGS. Finally and most critically, we found that including MPT in both MOGS and MOnGS calculations yields early learning scores that are significantly stronger predictors of the number of correct sequences performed by the end of acquisition. This indicates that ignoring MPT when calculating MOGS overlooks a critical aspect of the motor learning process. As a whole, our findings suggest that excluding MPT overinflates the magnitude of MOGS and decreases its relevance to overall motor learning.

Our core finding is that MOGS-TOTAL, which incorporates MPT, is negative rather than positive. This striking finding strongly suggests that traditional MOGS metrics, which do not account for MPT (i.e., MOGS-MET), inaccurately capture the true direction and magnitude of MOGS. Because KPS selectively measures motor execution speed, it artificially inflates performance by masking the time participants spend on motor planning. Importantly, Early learning-MET, which excludes motor preparation time, was not significantly correlated with the number of correct sequences performed. In contrast, we found that Early learning-TOTAL *was* significantly correlated with the number of correct sequences performed, indicating that including MPT in KPS calculation substantially increases the metric’s relevance to gold-standard learning metrics. This finding suggests that the traditional method for calculating MOGS and MOnGS does not comprehensively capture motor learning during ESL.

These results show that motor pre-planning is a key factor that contributes to performance changes across trials. Consistent with our findings, Das et al. (2025) recently reported that experimentally preventing participants from pre-planning the first few movements of an upcoming practice period significantly diminished MOGS magnitude. Building on these findings, our study quantified this pre-planning period using MPT and confirmed that including MPT in MOGS is necessary for accurately measuring motor learning. Because MOGS-MET measures motor execution only, it masks this motor planning delay. When this delay is factored in, MOGS-TOTAL becomes negative, indicating an actual *micro-offline loss* rather than a micro-offline gain.

Recent studies have questioned whether MOGS truly represent ‘rapid consolidation’. Instead, these studies suggest that MOGS merely reflect transient inter-trial performance changes, such as recovery from physical fatigue, rather than true offline memory stabilization (Ahmed et al., 2025; Das et al., 2025; Gupta & Rickard, 2024). Studies that calculated MOGS without MPT found that these early gains do not translate into long-term skill retention (Das et al., 2025). Furthermore, MOGS without MPT persists even when inter-trial rest periods are replaced with a cognitively engaging memory encoding task, indicating that true wakeful rest is not required for MOGS (Ahmed et al., 2025). Moreover, Das et al. (2025) specifically proposed and empirically demonstrated that motor pre-planning contributes to MOGS. Our study is the first to confirm these prior findings by directly quantifying MPT and also showing that it must be considered when developing valid learning metrics. Our findings are also consistent with performance-based explanations for MOGS, including reactive inhibition (Gupta & Rickard 2024). Finally, the dissociation between Early Learning-MET and the number correct sequences reported here supports the idea that MOGS-MET captures a temporary performance change rather than a persistent improvement reflective of learning.

Our study is not without limitations. A primary limitation of this study is that we pooled data from a study that originally had two groups. However, our results show that our manipulation (jittering the rest interval) was ineffective and did not produce meaningful group differences. Furthermore, without a random-sequence control group, this study cannot isolate whether MPT improvements represent sequence-specific skill acquisition or task-general motor execution gains. Future investigations should confirm these findings.

Overall, our results show that including MPT when calculating MOGS flipped it from positive to negative and improved its correlation with gold-standard ESL metrics. These findings suggest that including motor preparation time when calculating MOGS results in a more reliable and comprehensive metric of ESL. Additionally, our results show that MOGS are also a sensitive and volatile metric that should be interpreted with caution in motor learning research. Overall, results are consistent with the hypothesis that traditional MOGS calculations are driven by reductions in motor preparation time rather than learning or consolidation.

## Data availability

Datasets are available upon reasonable request.

## Author contributions

SJH and MF conceived and designed research; NA performed experiments; NA analyzed data; NA, TS, SJH and MF interpreted results of experiments; NA prepared figures; NA and MF drafted manuscript; NA, TS, SJH and MF edited and revised manuscript; NA, TS, SJH and MF approved final version of manuscript.

## References

Abrahamse, E., Ruitenberg, M., De Kleine, E., & Verwey, W. B. (2013). Control of automated behavior: Insights from the discrete sequence production task. Frontiers in Human Neuroscience, 7. 10.3389/fnhum.2013.00082

Ahmed, N. I., Suresh, T., Hussain, S. J., & Freedberg, M. (2025). Investigating the effect of task engagement during intertrial rest periods on micro offline gains. Scientific Reports, 15(1), 37396. 10.1038/s41598-025-21351-5

Albouy, G., Fogel, S., King, B. R., Laventure, S., Benali, H., Karni, A., Carrier, J., Robertson, E. M., & Doyon, J. (2015). Maintaining vs. enhancing motor sequence memories: Respective roles of striatal and hippocampal systems. NeuroImage, 108, 423–434. 10.1016/j.neuroimage.2014.12.049

Albouy, G., Sterpenich, V., Balteau, E., Vandewalle, G., Desseilles, M., Dang-Vu, T., Darsaud, A., Ruby, P., Luppi, P.-H., Degueldre, C., Peigneux, P., Luxen, A., & Maquet, P. (2008). Both the hippocampus and striatum are involved in consolidation of motor sequence memory. Neuron, 58(2), 261–272. 10.1016/j.neuron.2008.02.008

Ariani, G., & Diedrichsen, J. (2019). Sequence learning is driven by improvements in motor planning. Journal of Neurophysiology, 121(6), 2088–2100. 10.1152/jn.00041.2019

Ariani, G., Kwon, Y. H., & Diedrichsen, J. (2020). Repetita iuvant: Repetition facilitates online planning of sequential movements. Journal of Neurophysiology, 123(5), 1727–1738. 10.1152/jn.00054.2020

Bashford, L., Kobak, D., Diedrichsen, J., & Mehring, C. (2022). Motor skill learning decreases movement variability and increases planning horizon. Journal of Neurophysiology, 127(4), 995–1006. 10.1152/jn.00631.2020

Bönstrup, M., Iturrate, I., Hebart, M. N., Censor, N., & Cohen, L. G. (2020). Mechanisms of offline motor learning at a microscale of seconds in large-scale crowdsourced data. Npj Science of Learning, 5(1), 1–10. 10.1038/s41539-020-0066-9

Bönstrup, M., Iturrate, I., Thompson, R., Cruciani, G., Censor, N., & Cohen, L. G. (2019). A rapid form of offline consolidation in skill learning. Current Biology, 29(8), 1346–1351.e4. 10.1016/j.cub.2019.02.049

Brown, R. M., Friedgen, E., & Koch, I. (2022). The role of action effects in motor sequence planning and execution: Exploring the influence of temporal and spatial effect anticipation. Psychological Research, 86(4), 1078–1096. 10.1007/s00426-021-01525-2

Buch, E. R., Claudino, L., Quentin, R., Bönstrup, M., & Cohen, L. G. (2021). Consolidation of human skill linked to waking hippocampo-neocortical replay. Cell Reports, 35(10), 109193. 10.1016/j.celrep.2021.109193

Das, A., Karagiorgis, A., Diedrichsen, J., Stenner, M.-P., & Azañón, E. (2025). Micro-offline gains do not reflect offline learning during early motor skill acquisition in humans. Proceedings of the National Academy of Sciences, 122(44), e2509233122. 10.1073/pnas.2509233122

Diedrichsen, J., & Kornysheva, K. (2015). Motor skill learning between selection and execution. Trends in Cognitive Sciences, 19(4), 227–233. 10.1016/j.tics.2015.02.003

Fischer, S., Hallschmid, M., Elsner, A. L., & Born, J. (2002). Sleep forms memory for finger skills. Proceedings of the National Academy of Sciences, 99(18), 11987–11991. 10.1073/pnas.182178199

Gann, M. A., Dolfen, N., King, B. R., Robertson, E. M., & Albouy, G. (2023). Prefrontal stimulation as a tool to disrupt hippocampal and striatal reactivations underlying fast motor memory consolidation. Brain Stimulation, 16(5), 1336–1345. 10.1016/j.brs.2023.08.022

Gupta, M. W., & Rickard, T. C. (2024). Comparison of online, offline, and hybrid hypotheses of motor sequence learning using a quantitative model that incorporate reactive inhibition. Scientific Reports, 14(1), 4661. 10.1038/s41598-024-52726-9

Haith, A. M., Huberdeau, D. M., & Krakauer, J. W. (2015). The influence of movement preparation time on the expression of visuomotor learning and savings. Journal of Neuroscience, 35(13), 5109–5117. 10.1523/JNEUROSCI.3869-14.2015

Herszage, J., Sharon, H., & Censor, N. (2021). Reactivation-induced motor skill learning. Proceedings of the National Academy of Sciences, 118(23), e2102242118. 10.1073/pnas.2102242118

Hsu, G., Shereen, A. D., Cohen, L. G., & Parra, L. C. (2023). Robust enhancement of motor sequence learning with 4 mA transcranial electric stimulation. Brain Stimulation, 16(1), 56–67. 10.1016/j.brs.2022.12.011

Jacobacci, F., Armony, J. L., Yeffal, A., Lerner, G., Amaro, E., Jovicich, J., Doyon, J., & Della-Maggiore, V. (2020). Rapid hippocampal plasticity supports motor sequence learning. Proceedings of the National Academy of Sciences of the United States of America, 117(38), 23898–23903. 10.1073/pnas.2009576117

Karni, A., Meyer, G., Rey-Hipolito, C., Jezzard, P., Adams, M. M., Turner, R., & Ungerleider, L. G. (1998). The acquisition of skilled motor performance: Fast and slow experience-driven changes in primary motorJcortex. Proceedings of the National Academy of Sciences, 95(3), 861–868. 10.1073/pnas.95.3.861

Kogan, E., Lu, J., & Zuo, Y. (2023). Cortical circuit dynamics underlying motor skill learning: From rodents to humans. Frontiers in Molecular Neuroscience, 16. 10.3389/fnmol.2023.1292685

Krakauer, J. W., & Mazzoni, P. (2011). Human sensorimotor learning: Adaptation, skill, and beyond. Current Opinion in Neurobiology, Sensory and Motor Systems, 21(4), 636–644. 10.1016/j.conb.2011.06.012

Krakauer, J. W., & Shadmehr, R. (2006). Consolidation of motor memory. Trends in Neurosciences, 29(1), 58–64. 10.1016/j.tins.2005.10.003

Mylonas, D., Schapiro, A. C., Verfaellie, M., Baxter, B., Vangel, M., Stickgold, R., & Manoach, D. S. (2024). Maintenance of procedural motor memory across brief rest periods requires the hippocampus. Journal of Neuroscience, 44(14). 10.1523/JNEUROSCI.1839-23.2024

Nguyen, Q. N., Michon, K. J., Vesia, M., & Lee, T. G. (2025). Dissociable causal roles of dorsolateral prefrontal cortex and primary motor cortex over the course of motor skill development. Journal of Neuroscience, 45(20). 10.1523/JNEUROSCI.2015-23.2025

Nudo, R. J. (2013). Recovery after brain injury: Mechanisms and principles. Frontiers in Human Neuroscience, 7, 887. 10.3389/fnhum.2013.00887

Nudo, R. J., Wise, B. M., SiFuentes, F., & Milliken, G. W. (1996). Neural substrates for the effects of rehabilitative training on motor recovery after ischemic infarct. Science, 272(5269), 1791–1794. 10.1126/science.272.5269.1791

Psychology software tools, Inc. (2016). E-Prime [Computer software].

Robertson, E. M. (2007). The serial reaction time task: Implicit motor skill learning? The Journal of Neuroscience, 27(38), 10073–10075. 10.1523/JNEUROSCI.2747-07.2007

Robertson, E. M. (2012). New insights in human memory interference and consolidation. Current Biology, 22(2), R66–R71. 10.1016/j.cub.2011.11.051

Robertson, E. M., Pascual-Leone, A., & Press, D. Z. (2004). Awareness modifies the skill-learning benefits of sleep. Current Biology, 14(3), 208–212. 10.1016/j.cub.2004.01.027

Walker, M. P., Brakefield, T., Morgan, A., Hobson, J. A., & Stickgold, R. (2002). Practice with sleep makes perfect: Sleep-dependent motor skill learning. Neuron, 35(1), 205–211. 10.1016/S0896-6273(02)00746-8

Wang, Y., Huynh, A. T., Bao, S., Buchanan, J. J., Wright, D. L., & Lei, Y. (2024). Memory consolidation of sequence learning and dynamic adaptation during wakefulness. Cerebral Cortex, 34(2), bhad507. 10.1093/cercor/bhad507

Yang, Y., Huang, Z., Yang, Y., Fan, M., & Yin, D. (2025). Time-dependent consolidation mechanisms of durable memory in spaced learning. Communications Biology, 8(1), 535. 10.1038/s42003-025-07964-6

